# Rapid CRISPR–Cas9 Genome Editing in S. cerevisiae

**DOI:** 10.64898/2026.03.27.714888

**Authors:** Hosein Rostamian, Ethan W. Madden, Frank M. Kaplan, Riley Kim, Daniel G. Isom, Brian D. Strahl

**Author notes:** Technical contact (Hosein Rostamian).

## Abstract

This protocol enables rapid CRISPR-Cas9 genome editing in Saccharomyces cerevisiae by replacing restriction/ligation guide cloning with PCR-based protospacer installation and seamless plasmid recircularization. It describes in silico HDR donor and SgRNA design, install guide sequences into cas9 plasmid by PCR and seamless assembly, plasmid cloning and sequence verification in E. coli, and LiAc/PEG co-transformation of yeast with Cas9-sgRNA plasmid plus HDR donor. The workflow selects yeast colonies on G418 and confirms edits by PCR and sequencing.

## Before you begin

The CRISPR–Cas9 system enables programmable, sequence-specific genome editing ^1^. In *Saccharomyces cerevisiae*, a common bottleneck is the speed and reliability of constructing a Cas9 plasmid that carries the desired guide sequence. Many widely used yeast CRISPR plasmids rely on restriction enzyme–based guide cloning and ligation, which adds hands-on steps and introduces failure modes. These issues become especially limiting when generating multiple guides or iterating designs^2,3^.

This protocol describes a rapid workflow to introduce a user-defined single-guide RNA (sgRNA) into a Cas9 plasmid by whole-plasmid PCR, followed by seamless, homology-based DNA assembly to recircularize the plasmid^4^. To support this workflow, we generated an updated backbone, pML104-KanMX-sgRNAv2, which combines dominant KanMX/G418 selection with a redesigned guide-insertion site and an extended sgRNA scaffold. Together, these features support straightforward PCR-based protospacer replacement and compatibility with prototrophic strain backgrounds (e.g., CEN.PK). The complete workflow typically takes ∼10 days.

The steps below provide instruction for sgRNA selection, primer design to encode the sgRNA in the Cas9 plasmid, and HDR donor design for small-to-large deletions, point mutations, insertions/replacements, or tagging.

### Define the editing objective

Timing: 10–20 min

1. Decide whether you will perform a deletion, point mutation, insertion/replacement, or N- or C-terminal tagging.
2. Obtain the genomic sequence and annotation needed for gene targeting (e.g., from SGD or your lab’s strain reference).

**Note:** Most reference genomic information is derived from the S288C strain. If your experiments use a different genetic background or yeast species, confirm the gene sequence and genomic context using strain-specific resources or sequence the target region.

### Design guide RNA

Timing: 30-60 min

The CRISPR–Cas9 system uses a sgRNA to direct *Streptococcus pyogenes* Cas9 (SpCas9) to a specific genomic site. SpCas9 binding and cleavage require two sequence features: (i) a protospacer-adjacent motif (PAM) in the genome, and (ii) a neighboring 20-nt protospacer immediately upstream of the PAM. In this protocol, the “guide sequence” refers to the 20-nt protospacer sequence (written 5ʹ→3ʹ) that is encoded within the sgRNA and determines target specificity.

**3.** Apply SpCas9 guide-design rules.

a. **PAM requirement**: Identify candidate sites with a 5ʹ-NGG-3ʹ PAM in the genome.
b. **Guide definition**: Define the guide as the 20 nt immediately 5ʹ of the PAM on the PAM-containing strand.
c. **Expected cut site**: SpCas9 typically cleaves ∼3 bp upstream of the PAM (toward the 5ʹ end of the protospacer).
d. **Do not include the PAM**: Do not clone or order the NGG PAM as part of the guide sequence.
e. **Either strand is valid**: Candidate guides can be selected from NGG sites on either strand.

Example: For the genomic sequence (5ʹ→3ʹ): 5ʹ-GATTCCAG TACGCATCGGTGGACTATCG ***CGG*** TAGTATC -3ʹ; the PAM is 5ʹ-CGG-3ʹ and the corresponding guide sequence (20-nt protospacer) is 5ʹ-TACGCATCGGTGGACTATCG-3ʹ.

**4.** Select candidate sgRNAs

a. Identify candidate sgRNAs using a computational guide-design tool (e.g., Benchling).
b. Log in to Benchling (benchling.com) and start a new guide design**: Create → CRISPR → CRISPR Guide**.
c. Enter the target gene name/ID (or paste the target sequence), then confirm the organism and target region displayed by Benchling.
d. If designing for N- or C-terminal tagging, open **Advanced settings** and extend the target region upstream and/or downstream of the gene to include the start or stop codon region.
e. Click **Next**, review settings (default parameters are typically sufficient), then click **Finish** to generate candidate guides.
f. In the results view, select the sequence window where you want cleavage and click **Add (+)** to generate guides for that region (Figure 1).
g. Rank candidates by predicted on-target score and review predicted off-targets, then select 2–3 sgRNAs for downstream cloning.

**Figure 1.**
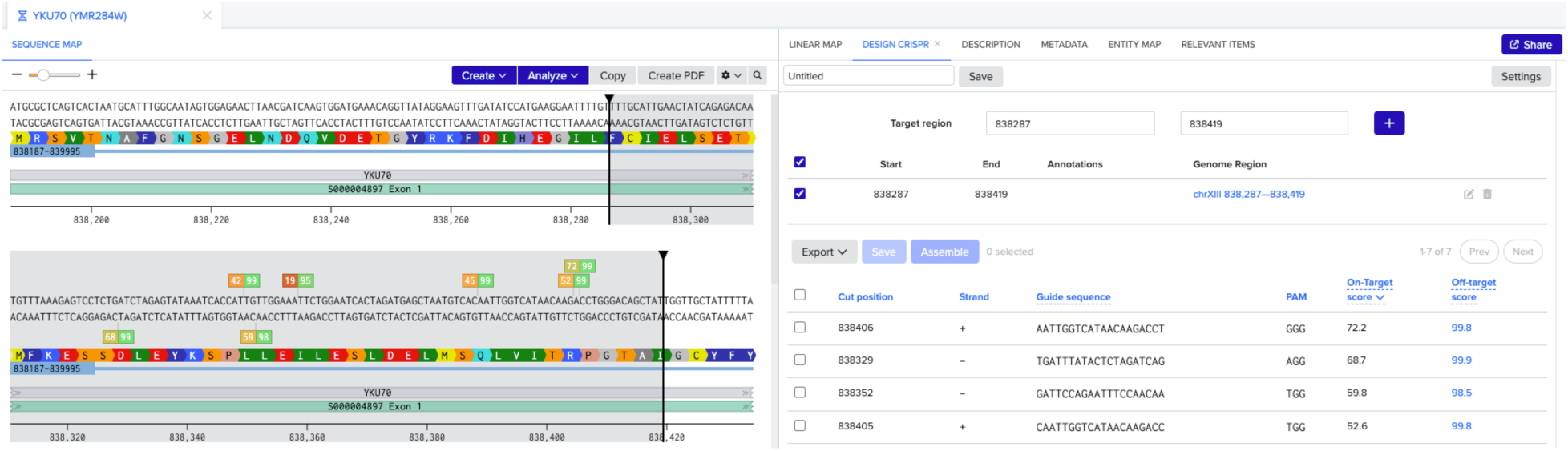
Designing sgRNAs in Benchling for *S. cerevisiae*.(A) Benchling sequence view of the target locus showing the selected region used for guide design. **(B)** Benchling CRISPR Guide results table listing candidate sgRNAs with cut position/strand, guide sequence and PAM, and predicted on-target and oD-target scores used to rank and select guides.

**Note:** Practical selection criteria for sgRNAs include high predicted on-target activity, low predicted off-target risk, and avoidance of guides with extreme GC content or long homopolymer runs.

**Note**: Design and carry forward at least two independent sgRNAs per target locus, as editing efficiency can vary between guides and one may fail or perform poorly.

**5.** Choose sgRNA placement based on the edit type.

a. **Deletion**: Select an sgRNA within the region to be deleted.
b. **Point mutation or small insertion**: Select an sgRNA as close as possible to the intended edit site; when feasible, within ∼100 bp.
c. **Protein N-terminal tagging**: Select an sgRNA within ∼100 bp upstream or downstream of the start codon.
d. **Protein C-terminal tagging**: Select an sgRNA within ∼100 bp upstream or downstream of the stop codon.

Note: While expanding the search region can increase the number of candidate sgRNAs, edits that rely on HDR (e.g., point mutations and tagging) are generally most efficient when cleavage occurs near the desired modification site.

### Design primers to incorporate the sgRNA into the Cas9 plasmid

Timing: 20 min

In this section, design a forward and reverse primer pair to introduce the 20-nt guide (protospacer) sequence into the Cas9 plasmid by whole-plasmid PCR. Each primer contains (1) a guide-specific segment and (2) a constant vector-homology tail. The guide-specific segments are in opposite orientations so that the resulting amplicon encodes the desired sgRNA sequence (see Figure 2).

**6.** Design primers to introduce the 20-nt guide sequence by PCR.

a. Identify the protospacer insertion site in the Cas9 plasmid backbone (see Figure 2).
b. In SnapGene, open the pML104-KanMX-sgRNAv2 plasmid and replace the existing 20-nt protospacer with your selected 20-nt guide sequence from the previous section.
c. Design a **59-nt forward primer** that includes:

i. a **17-nt guide-specific segment** derived from the 20-nt protospacer.
ii. a **42-nt constant vector-homology segment** that anneals to the plasmid backbone.
d. Design a **50-nt reverse primer** that includes:

i. an **18-nt guide-specific segment** derived from the 20-nt protospacer.
ii. a **32-nt constant vector-homology segment** that anneals to the plasmid backbone.
e. Ensure the two primers create a 15bp 5ʹ overlap (typically 15–20 bp) to enable seamless recircularization of the PCR product during assembly.

**Figure 2.**
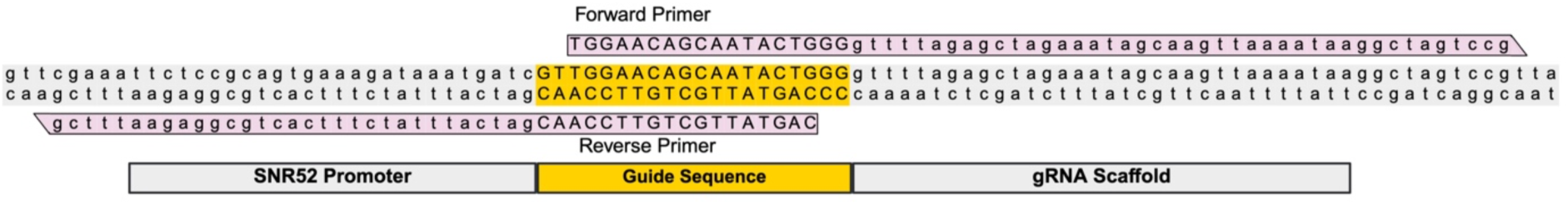
**Primer Design for PCR-based sgRNA insertion into pML104-KanMX-sgRNAv2 Vector**: Schematic of forward (F) and reverse (R) primer design for the insertion of sgRNA into the pML104-KanMX-sgRNAv2 vector using the In-Fusion Snap Assembly Master Mix. The yellow box represent the guide sequence within the sgRNA cassette of the vector. The forward primer (F) includes 17 nucleotides of the sgRNA sequence and 42 nucleotides of the vector sequence, while the reverse primer (R) includes 32 nucleotides of the vector sequence and 18 nucleotides of the sgRNA sequence. The primers have a 15-base pair overlap, enabling seamless assembly of the linearized vector without the use of restriction enzymes or ligase.

CRITICAL: Primer length and overlap strongly affect plasmid amplification and assembly. The primer lengths and homology tails listed here were optimized for the pML104-KanMX-sgRNAv2 backbone and In-Fusion Snap Assembly Master Mix (Takara). If using a different plasmid or assembly system, keep the design strategy but adjust overlap length and homology tails according to the manufacturer’s recommendations.

### Design homology-directed repair (HDR) template

Timing: 30-60 min

The HDR donor contains the desired modification flanked by left and right homology arms matching the target locus, enabling precise integration by Homology-directed repair. In this protocol, we use vendor-synthesized dsDNA fragments (e.g., Twist) as HDR donors because they are cost-effective, support complex edits (e.g., tag plus point mutation) in a single construct.

Optional: HDR donors can also be generated by no-template overlap/assembly PCR instead of ordering synthetic dsDNA; primer-design guidance is available^5^.

**7.** Define the HDR donor architecture for the intended edit.

**a. Design HDR template for gene or sequence deletion**

i. Determine the genomic location of the target gene or sequence to be deleted.
ii. Select the sequences flanking the region to be deleted.
iii. Define the **left homology arm** as ∼200-300 bp upstream of the deletion boundary. For gene deletions, this corresponds to the upstream of the ATG start codon.
iv. Define the **right homology arm** as ∼200-300 bp downstream of the deletion boundary. For gene deletions, this corresponds to downstream of the stop codon.
v. Assemble the HDR donor so the left and right homology arms are **directly adjacent,**
vi. mitting the deleted region (Figure 3).
vii. Introduce silent mutations in the PAM and/or guide-binding region to prevent re-cutting (see next step).
**b. Design HDR template for point mutation**

i. Determine the specific genomic position where the point mutation is desired and identify a nearby sgRNA/PAM site.
ii. Select DNA sequences flanking the intended edit site, typically ranging from ∼200-300 bp on each side, to serve as left and right homology arms.
iii. Replace the original codon(s) with the new codon(s) corresponding to the desired amino acid substitution. For example, if the goal is to change an Glutamic acid residue to alanine, the DNA codons for isoleucine “GAA” would be replaced with the codons for alanine “GCA” **(Figure 4)**.
iv. Introduce silent mutations in the PAM and/or guide-binding region to prevent re-cutting (see next step).
**c. Design the HDR donor for N-terminal tagging**

i. Identify the start codon and confirm the tag sequence will be in-frame with the coding sequence (Figure 5).
ii. Build the donor so the tag is inserted immediately after the ATG.
iii. Include a left homology arm upstream of the ATG (typically ∼200-300 bp).
iv. Include the ATG in the donor sequence.
v. Place the tag sequence directly after the ATG and maintain the reading frame.
vi. Include a right homology arm downstream of the insertion junction (typically ∼200-300 bp).
vii. Introduce silent mutations in the PAM and/or guide-binding region to prevent re-cutting (see next step).
**d. Design the HDR donor for C-terminal tagging**

i. Identify the stop codon and confirm the tag sequence will be in-frame with the coding sequence (Figure 5).
ii. Build the donor so the tag is inserted immediately before the stop codon.
iii. Include a left homology arm upstream of the stop codon (typically ∼200-300 bp).
iv. Place the tag sequence directly before the stop codon and maintain the reading frame.
v. Include the stop codon in the donor sequence.
vi. Include a right homology arm downstream of the stop codon (typically ∼200--300 bp).
vii. Introduce silent mutations in the PAM and/or guide-binding region to prevent re-cutting (see next step).

**Figure 3.**
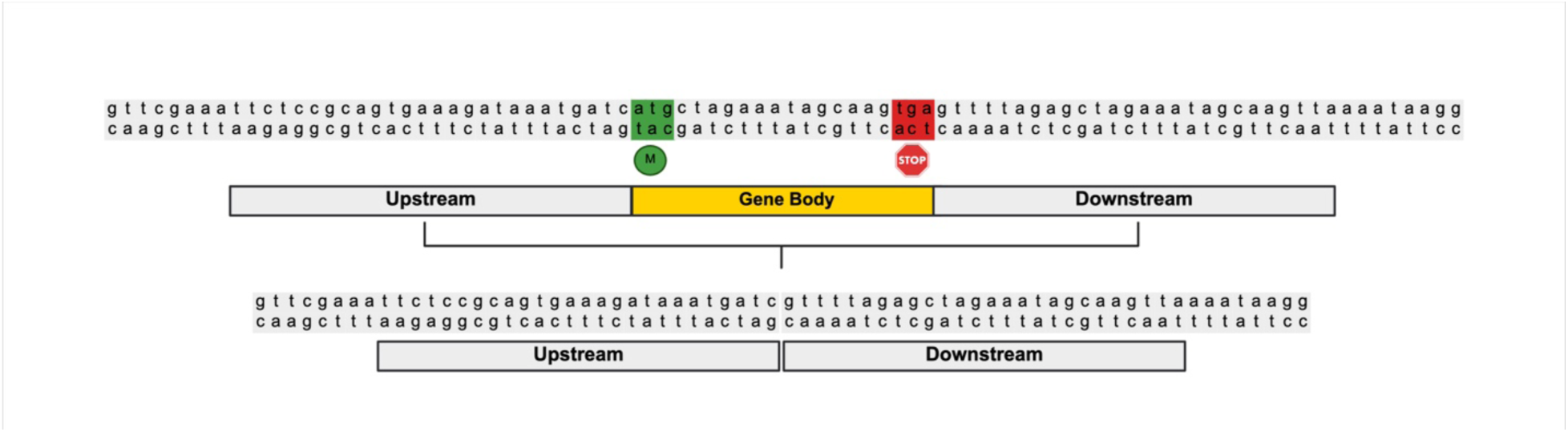
HDR donor design for gene/sequence deletion. **(A)** Schematic showing endogenous locus targeted for deletion (coding sequence between the ATG start codon and STOP codon). The upstream and downstream flanking sequences are selected as homology arms that border the deletion boundaries. **(B)** HDR donor used for deletion. The donor consists of the upstream homology arm directly fused to the downstream homology arm, with no intervening target sequence. After Cas9 cutting, homology-directed repair uses this donor to join the upstream and downstream regions, resulting in precise deletion of the target gene/sequence.

**Figure 4.**
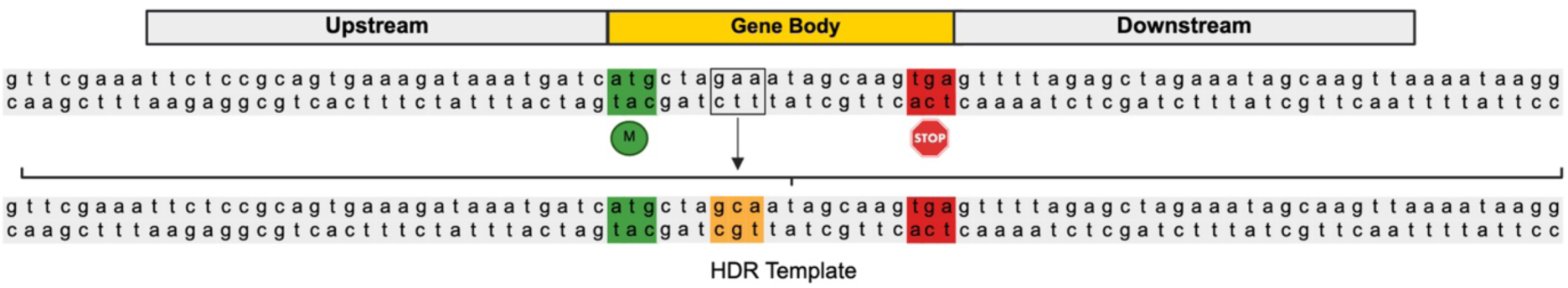
**Design of HDR Template for Point Mutation**: Schematic representation of a HDR template designed for introducing a point mutation. The HDR template includes a mutation from Glutamic acid to Alanine (Glu to Ala).

**Figure 5.**
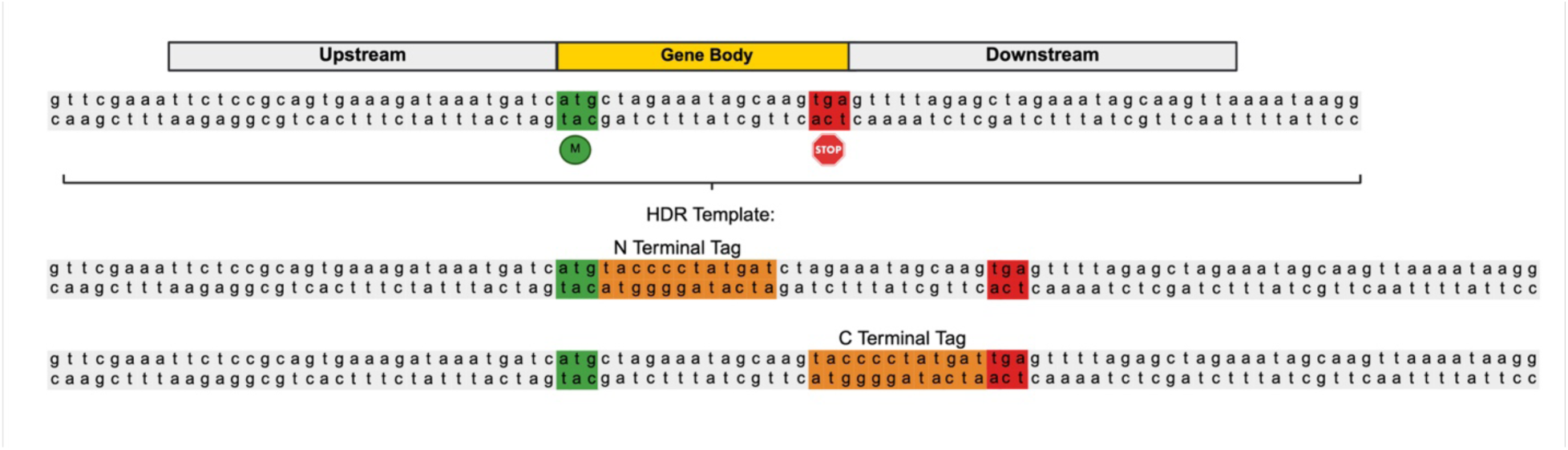
HDR donor design for epitope tagging. **(A) N-terminal tagging strategy.** Schematic of a genomic locus showing the ATG start codon and the selected upstream and downstream homology arms (typically 200–300 nt each) flanking the start site. The tag sequence is inserted immediately after the ATG in the HDR donor and kept in-frame with the coding sequence. **(B) C-terminal tagging strategy.** Schematic showing the stop codon and the upstream and downstream homology arms (typically 200–300 nt each) flanking the stop site. The tag sequence is inserted immediately before the stop codon in the HDR donor, followed by the stop codon and downstream homology arm.

Note: Some epitope tags (e.g., 3×HA) are highly repetitive at the DNA level, and some vendors will not synthesize HDR templates containing long repeats. To avoid this, recode the tag with synonymous codons to reduce nucleotide repetition while keeping the amino acid sequence unchanged (e.g., alternate alanine codons: GCT → GCC/GCA).

**e. Design HDR template for insertion or replacement**

i. Identify the intended integration site (or replacement boundaries) in the genome.
ii. Define the left and right homology arms flanking the insertion/replacement site (typically ∼200–300 bp on each side).
iii. Insert the desired sequence between the homology arms in the HDR template.
iv. Introduce silent mutations in the PAM and/or guide-binding region to prevent re-cutting (see next step).

Note: If the insertion is within a coding sequence, design the donor so the insert is **in-frame** and does not introduce unintended stop codons or frameshifts.

Note: Check the final edited sequence to ensure the insert or junctions do not create new Cas9 target sites (e.g., a new NGG PAM adjacent to a matching protospacer) near the edited locus.

### Prevent Cas9 re-cutting by mutating PAM/protospacer

Timing: 10–20 min

After HDR, the edited locus can remain a valid Cas9 target if the PAM and protospacer sequence are unchanged. Continued Cas9 cutting at the repaired locus can reduce recovery of correctly edited clones. To prevent re-cutting, introduce mutations in the HDR donor that disrupt Cas9 recognition while preserving the intended final edited sequence. For coding regions, introduce **silent mutations**, so the protein sequence is unchanged.

**8.** Introduce protective mutations in the HDR donor to prevent Cas9 re-cutting

a. Choose one (or both) of the following protection strategies

i. **Mutating the PAM site**: Introduce a substitution within the NGG PAM so it no longer matches SpCas9 requirements.

Example: If the original PAM site is 5’-CGG-3’ (coding for arginine), mutate it to a synonymous arginine codon such as AGA to disrupt the PAM while preventing the amino acid sequence **(Figure 6)**.

ii. **Mutating the sgRNA binding site(protospacer)**: Introduce ≥2 substitutions within the ∼10 bp region immediately upstream of the PAM (the “seed region”) to disrupt sgRNA binding. **(Figure 6)**.

**Figure 6.**
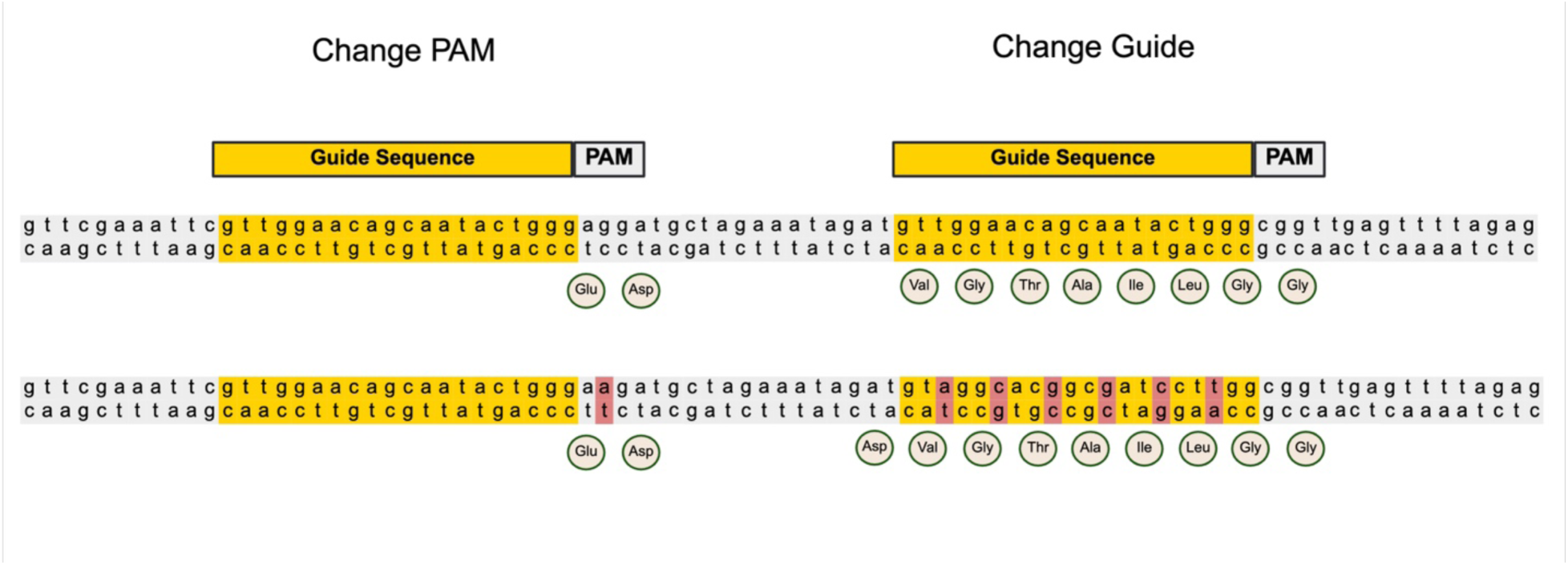
Design of an HDR template incorporating silent mutations to prevent Cas9 re-cutting: The schematic illustrates silent mutations introduced into the PAM site of guide seuqnce1 and the binding site of guide seuqnce2 to block Cas9 recognition following HDR integration. Nucleotide substitutions are shown in red.

Example: For an sgRNA target sequence such as 5ʹ-ATT GTA TTG TCA CCG ACT ACA-3ʹ, introduce ≥2 synonymous substitutions within the ∼10 bp seed region to reduce sgRNA binding (e.g., TGT→TGC [Cys], ATT→ATA [Ile], GTC→GTA [Val], ACC→ACA [Thr]) without altering the protein sequence.

**CRITICAL**: When disrupting the PAM or protospacer, verify that the edited sequence does **not** introduce a new nearby PAM (e.g., NGG or other permissive PAMs such as NAG) that could allow continued Cas9 targeting.

### Design primers for HDR template amplification

Timing: 10–20 min

**9.** Design primers to PCR-amplify the HDR donor at the scale needed for transformation.

a. If using a vendor-synthesized dsDNA donor, design a **forward primer** that anneals at the **5ʹ end** of the donor and a **reverse primer** that anneals at the **3ʹ end** of the donor to amplify the full-length HDR template (Figure 7).
b. Plan amplification to generate sufficient donor DNA for yeast transformation, **typically ≥10 µg total donor DNA**.

**Figure 7.**
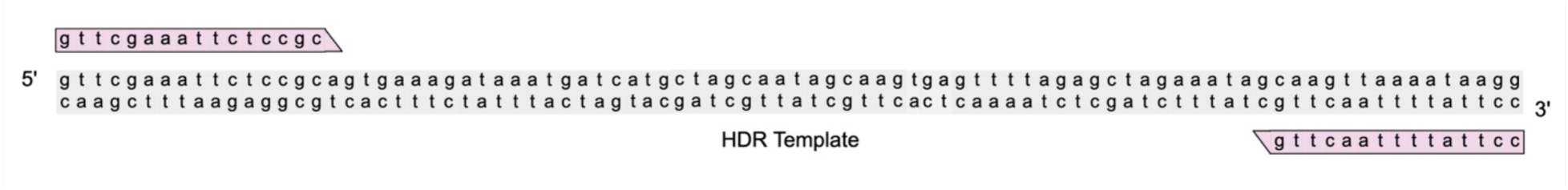
Primers for PCR amplification of HDR templates. Forward and reverse primers anneal to the 5′ and 3′ ends of the HDR donor to amplify the full-length template and generate suDicient DNA for yeast transformation.

Note: Vendor-supplied dsDNA fragments are usually delivered at amounts below what is required for yeast transformation; PCR amplification is used here to increase donor yield.

### Design primers to genotype CRISPR-edited yeast

This section describes how to design PCR primers that distinguish correctly edited clones from unedited cells after CRISPR-Cas9-mediated genome editing with an HDR donor.

**10.** Design PCR primers that flank the intended edit.

a. Place the forward primer upstream of the left homology arm (i.e., outside the HDR donor sequence).
b. Place the reverse primer downstream of the right homology arm (i.e., outside the HDR donor sequence).
c. Ensure the amplicon spans the edit junction(s) so that edited and unedited alleles yield different-size products and/or can be distinguished by sequencing.
d. Verify primer specificity in silico against the yeast genome and avoid primers that anneal within repeated elements.

Note: For deletions or insertions, design primers so the edited allele produces a clear size shift relative to an unedited control. For point mutations, the PCR product size may be unchanged, so plan to confirm the edit by sequencing across the modified region.

### Obtain chemically competent E. coli cells

**11.** Obtain chemically competent *E. coli* cells for plasmid cloning.

a. **Commercial option**: Use commercially prepared, high-efficiency chemically competent cells and follow the manufacturer’s transformation instructions.
b. **In-house option**: Prepare chemically competent cells using a standard calcium chloride/heat-shock protocol^6^.

Note: Use a high-efficiency cloning strain (e.g., DH5α) to maximize recovery of assembled plasmids.

### Prepare plates and media for selection

Timing: 2–6 h

**12.** Prepare LB agar plates for *E. coli* using a standard LB/LA recipe^7^.

a. Autoclave LB agar and cool to ∼50–55°C,
b. add kanamycin to the working concentration used for your plasmid selection (commonly 50 µg/mL), mix thoroughly, then pour plates.
**13.** Prepare LB liquid medium for *E. coli* using a standard LB recipe^7^.

a. After sterilization and cooling, add kanamycin to the working concentration (commonly 50 µg/mL).
**14.** Prepare YPD liquid medium for yeast using a standard YPD recipe^8^.
**15.** Prepare YPD agar plates containing G418 for yeast selection using a standard YPD agar recipe.

a. Autoclave YPD agar and cool to ∼50–60°C
b. Add G418 (commonly 0.5 mg/mL), mix thoroughly, then pour plates.

## Innovation

This protocol streamlines CRISPR–Cas9 editing in *Saccharomyces cerevisiae* by simplifying guide construction and standardizing HDR donor preparation. Guides are installed by PCR-based protospacer replacement using guide-encoding primers, followed by a single seamless-assembly step. This avoids restriction/ligation cloning, reduces common cloning failures, and rapid guide iteration. We also introduce a KanMX/G418-selectable Cas9–sgRNA backbone (pML104-KanMX-sgRNAv2) with a redesigned guide insertion site and an extended sgRNA scaffold. These features support straightforward PCR-based protospacer replacement and facilitate CRISPR editing in *S. cerevisiae*, including prototrophic strains such as CEN.PK. HDR donors are standardized using vendor-synthesized dsDNA templates with consistent homology-arm and PAM/seed-disruption rules. This supports deletions, point mutations, insertions, tagging, and combined edits (e.g., tag plus point mutation) in a single construct while reducing variability and PCR-related donor failures. Together, these updates provide a compact, teachable workflow for faster, more reproducible design-to-validated-edit cycles.

## Institutional permissions

Not applicable (work in *Saccharomyces cerevisiae*).

### Key resources table

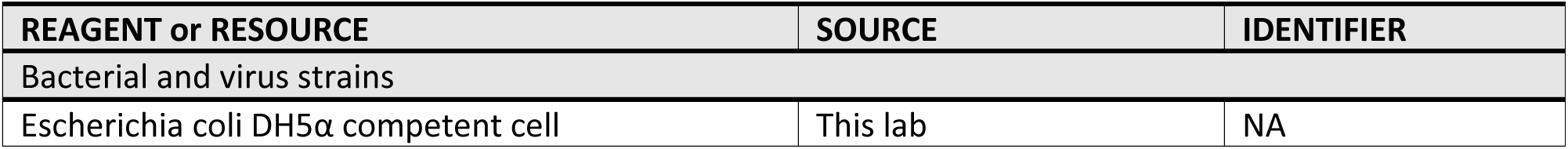
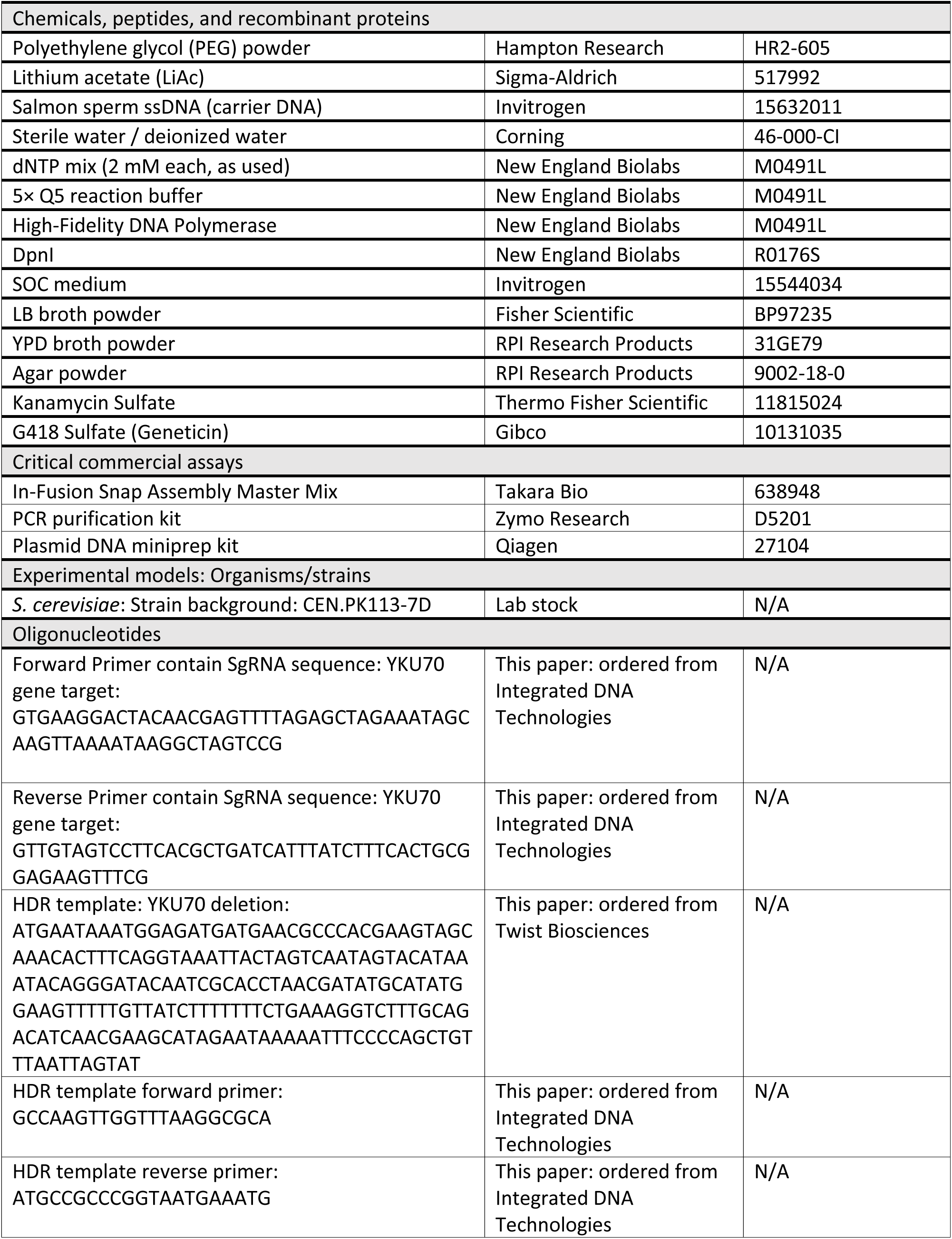
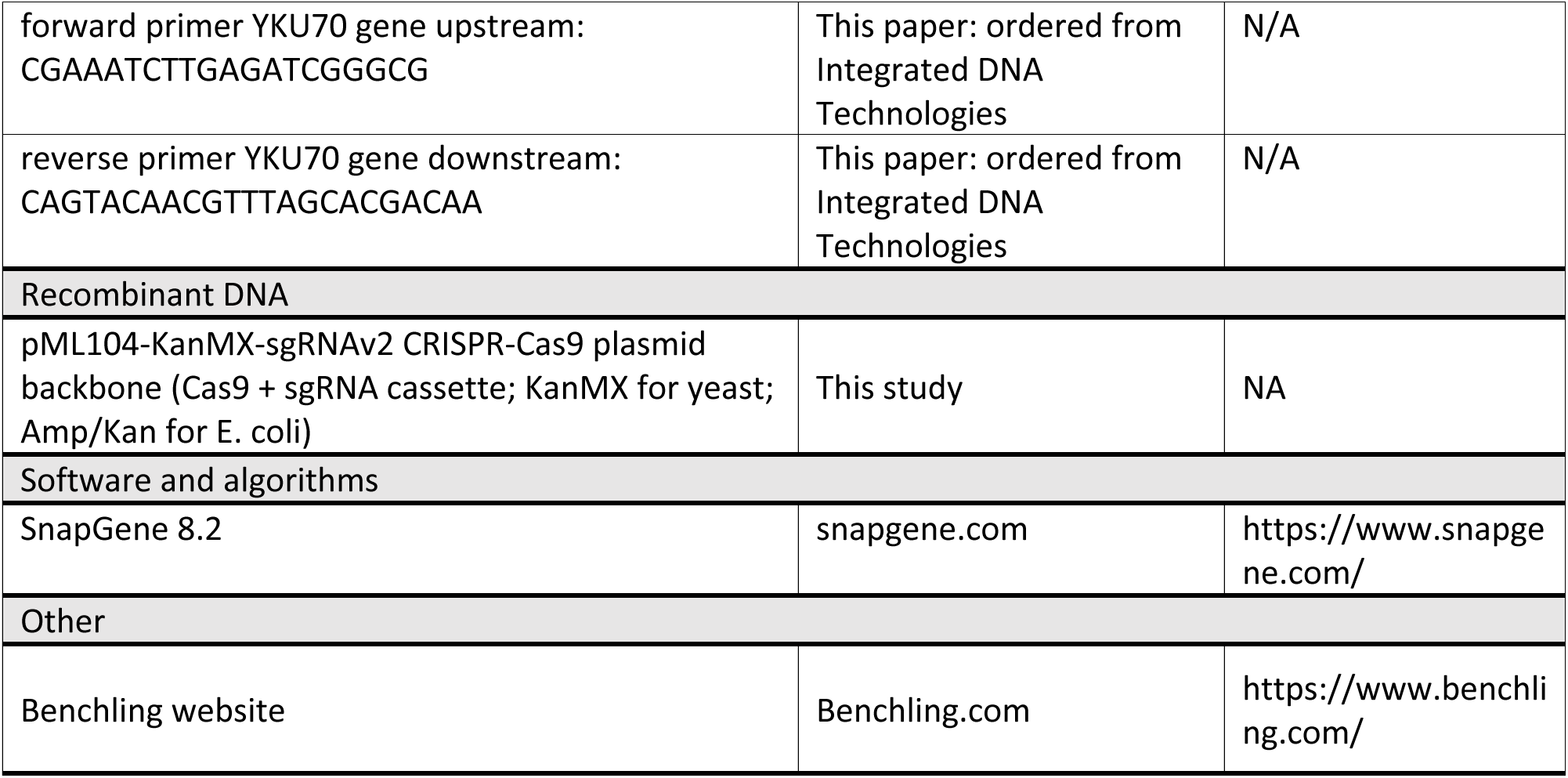

### Step-by-step method details

#### PCR-based sgRNA insertion into the CRISPR plasmid

Timing: 7 h

This major step amplifies the full CRISPR plasmid by whole-plasmid PCR while replacing the 20-nt guide sequence using primers that carry the target-specific sgRNA. At the end of this step, you will have a full-length PCR product (≈12 kb) containing the desired sgRNA sequence for downstream processing.

Note: This protocol uses the pML104-KanMX-sgRNAv2 Cas9 plasmid (≈12 kb). During PCR, the plasmid’s 20-nt guide region is swapped for a new user-defined sgRNA guide sequence. The plasmid is propagated in *E. coli* (Amp/Kan) and maintained in yeast by KanMX selection on G418.

1. Assemble a 50 μL PCR reaction on ice.

a. Add the following components to a 0.2 mL PCR tube for one rection (scale as needed):

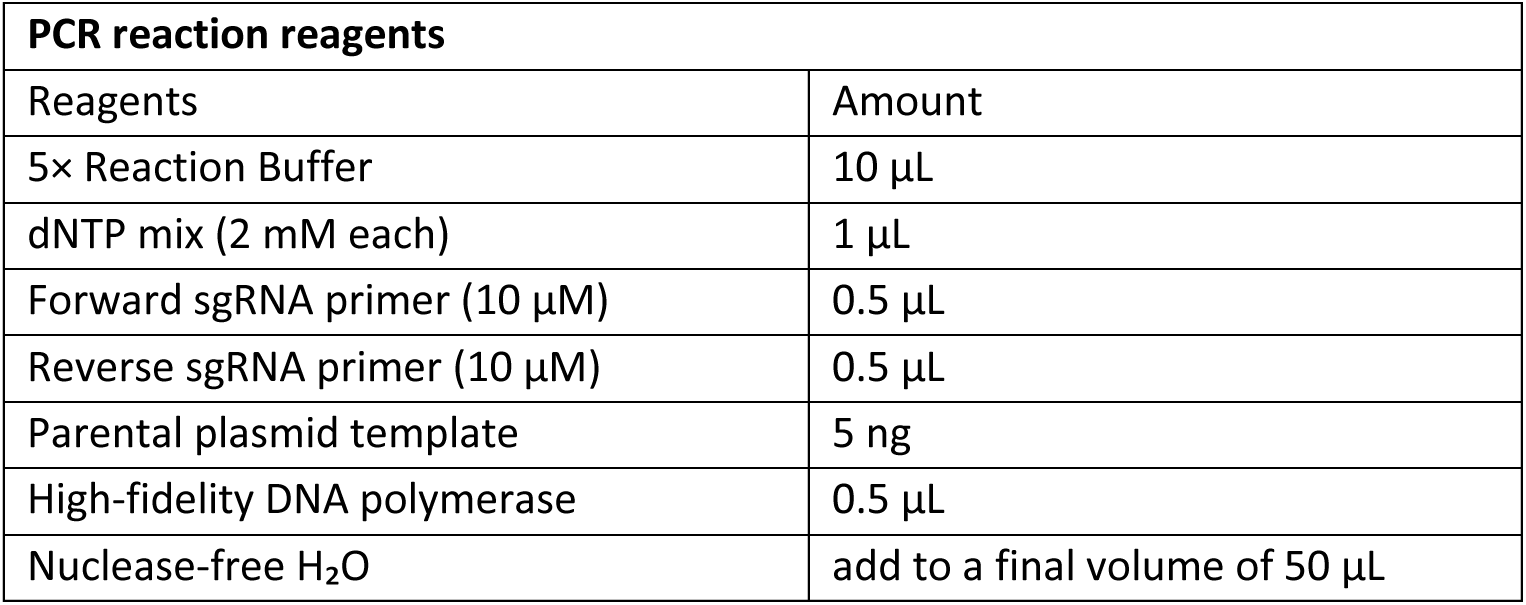

b. Mix gently by pipetting, then briefly spin down the reaction.

CRITICAL: Use a low primer concentration; excess primer can increase non-specific amplification and generate incorrect-sized products.

Note: Use a high-fidelity, proofreading polymerase (e.g., Q5 from New England Biolabs) to minimize point mutations that can reduce editing efficiency or plasmid function.

2. Run PCR using a gradient annealing temperature.

a. Place the tubes in a thermocycler
b. Program the annealing step as an 8-point gradient from 48°C to 63°C to identify the optimal annealing temperature
c. Run the PCR program as follows (Optionally overnight)

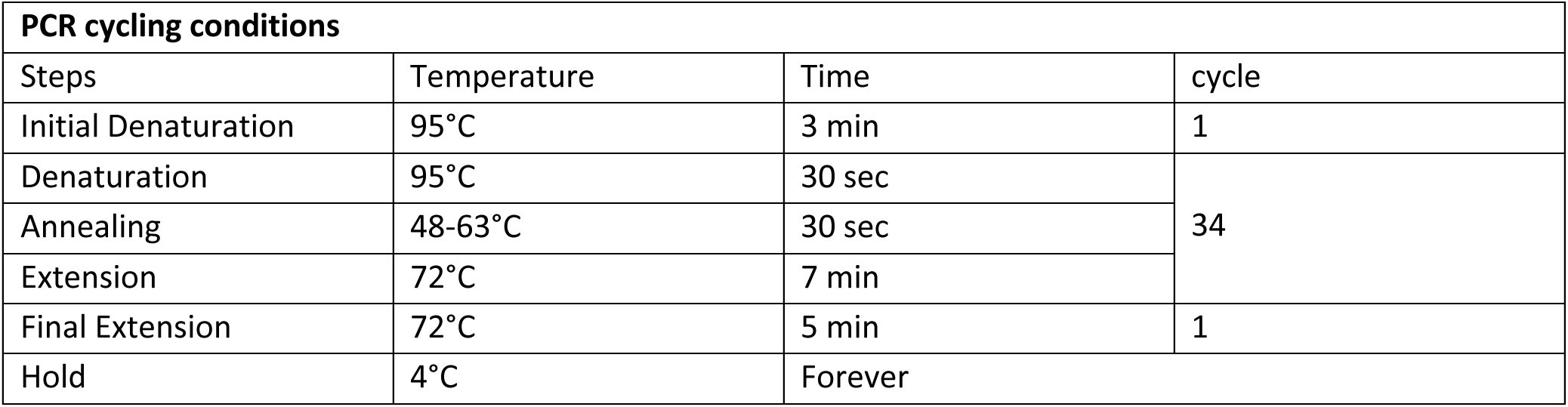

| PCR cycling conditions |  |  |  |
| --- | --- | --- | --- |
| Steps | Temperature | Time | cycle |
| Initial Denaturation | 95°C | 3 min | 1 |
| Denaturation | 95°C | 30 sec | 34 |
| Annealing | 48-63°C | 30 sec |  |
| Extension | 72°C | 7 min |  |
| Final Extension | 72°C | 5 min | 1 |
| Hold | 4°C | Forever |  |

3. Verify amplification by agarose gel electrophoresis.

a. Load 10 μL of the PCR reaction on a 0.7%–1% agarose gel with an appropriate DNA ladder.
b. Confirm a strong band at the expected full-length size (≈12 kb).
c. If additional non-specific bands are present, proceed if the correct-sized band is strong relative to other products (see Figure 8).
d. Store the PCR product until after plasmid sequence verification after *E. coli* transformation. If cloning or assembly fails on the first attempt, reuse the PCR product.

**Figure 8:**
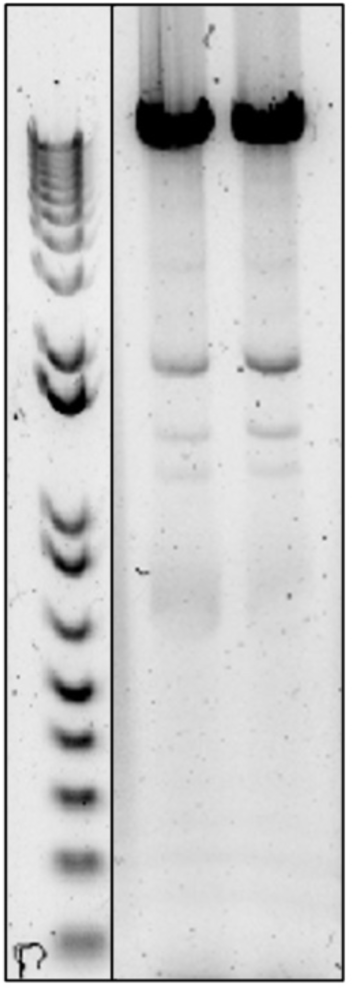
Agarose gel electrophoresis results of PCR products for inserting sgRNA sequences into the CRISPR vector backbone. The expected-size band in each sample lane indicates successful amplification of the linearized vector backbone with incorporation of the target sgRNA sequence. Additional faint bands represent non-specific amplification products.

#### DpnI digestion and PCR clean up

Timing: 90 min

This step digests the methylated parental plasmid template with DpnI while leaving the unmethylated PCR-amplified plasmid intact and then purifies the product for downstream transformation.

4. Add 2 μL of DpnI directly to the PCR product, mix by pipetting, and briefly spin down.
5. Place the tube in a thermocycler.
6. Incubate at 37°C for 60 min.
7. Heat-inactivate DpnI at 80°C for 20 min.
8. Purify the digested PCR product using a PCR purification kit according to the manufacturer’s instructions.

#### Seamless Circularization of the CRISPR plasmid

Timing: 20 min

This step circularizes the PCR-amplified CRISPR plasmid backbone using In-Fusion Snap Assembly Master Mix (Takara Bio) to generate a transformable plasmid.

9. Assemble a 10 μL In-Fusion Snap Assembly reaction.

a. Add 2 μL of 5x In-Fusion Snap Assembly Master Mix.
b. Add 200 ng of PCR-amplified vector
c. Add nuclease-free water to a final volume of 10 μL.
10. Mix by pipetting and briefly spin down.
11. Incubate at 50°C for 15 min in a thermocycler.
12. Place the reaction on ice.
13. Store the reaction at −20°C until bacterial transformation.

#### Transform CRISPR plasmid into DH5α E. coli

Timing: 3 days

This step amplifies the circularized CRISPR plasmid and generates enough high-quality plasmid DNA for sequence verification and downstream yeast transformation. Transformants are selected on kanamycin to maintain the plasmid during propagation.

14. Place the competent DH5a *E. coli* cells on ice to thaw.
15. Gently mix and then transfer 50 µl into a 1.5 ml Eppendorf tube. Avoid using a vortex.
16. Add 5 µl of the In-Fusion reaction mixture to the cells. Do not exceed 5 µl as it can inhibit transformation.
17. Incubate the tubes on ice for 30 min.
18. Place cells in a water bath set to 42°C and incubate for exactly 45 seconds.
19. Immediately return the tubes to ice for 2 minutes
20. Add pre-warmed (37°C) SOC or LB media to a final volume of 500 µl.
21. Incubate the cells at 37°C for 1 hour with shaking at 160–225 rpm.
22. Centrifuge at 3800g for 5 min, remove the supernatant, and resuspend the pellet in 100 µL SOC.
23. Spread the resuspended cells evenly onto the LB agar with kanamycin plates.
24. Incubate plates overnight at 37°C.
25. Next day Pick 4 well-isolated colonies and inoculate each into 5 mL LB containing kanamycin.
26. Grow cultures overnight at 37 °C with shaking (≈200–250 rpm).
27. Isolate plasmid DNA from at least 2 cultures using a standard miniprep kit.
28. Quantify plasmid using a Nanodrop.
29. Verify correct sgRNA insertion by sequencing

#### Amplify HDR template

Timing: 3–5 h total (hands-on: 45–60 min)

This step PCR-amplifies the HDR template to generate sufficient DNA for yeast transformation.

30. Set up a 50 µL PCR using 10 ng of the HDR fragment.

a. Use primers complementary to the 5’ and 3’ ends of the HDR fragment.
b. Run 6 parallel PCR reactions per HDR template to generate sufficient DNA for multiple yeast transformations.

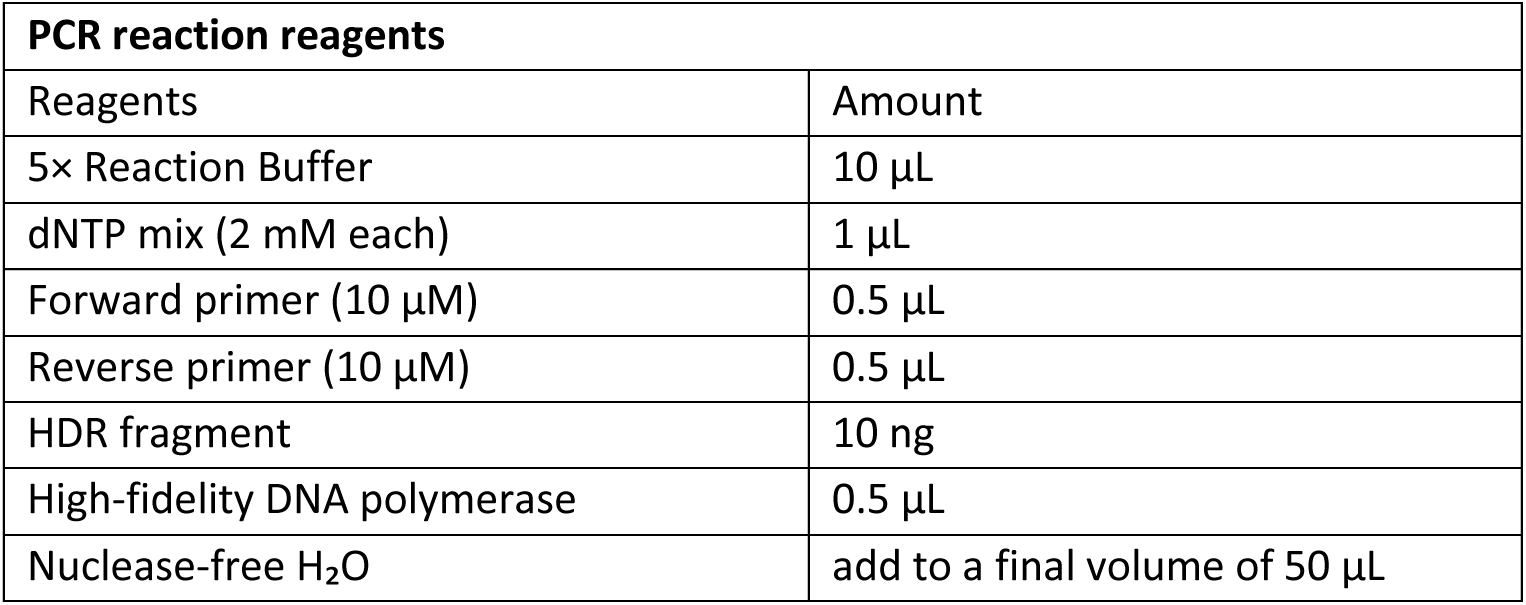

31. Run the PCR program as follows

| PCR cycling conditions |  |  |  |
| --- | --- | --- | --- |
| Steps | Temperature | Time | cycle |
| Initial Denaturation | 95°C | 3 min | 1 |
| Denaturation | 95°C | 30 sec | 34 |
| Annealing | 48-63°C | 30 sec |  |
| Extension | 72°C | varied<br>(30 second per 1Kb size of the product) |  |
| Final Extension | 72°C | 5 min | 1 |
| Hold | 4°C | Forever |  |

32. Pool the PCR reactions into a single tube.
33. Quantify HDR template DNA using a NanoDrop.
34. Verify HDR template size by running 10 μL of the purified product on an agarose gel and confirm a single band at the expected size.

### Transform yeast with CRISPR plasmid + HDR template

Timing: 4 days

This step introduces the Cas9/sgRNA plasmid and HDR template into yeast using a high-efficiency lithium acetate (LiAc)/ polyethylene glycol (PEG) transformation^9^ and then selects for transformants on G418.

CRITICAL: Throughout this procedure, use sterile tubes, tips, and solutions, and perform all steps aseptically near a flame to reduce contamination risk.

35. Grow an overnight yeast culture in YPD to reach log phase.

a. Calculate the culture volume needed for the planned number of transformations and include one additional transformation for the no-repair (–HDR) control.
b. Inoculate a yeast colony in 5 mL of YPD per planned transformation (e.g., 15 mL YPD for 3 transformations).
c. Incubate at 30°C with shaking (220 rpm) until the culture reaches (OD_600_ = 0.6-1).
d. If the culture is over log phase the next morning, dilute to OD_600_ to 0.2-0.4 in fresh YPD and continue incubating at 30°C until it returns to log phase (OD_600_= 0.6-1).
36. Harvest cells for each transformation and for the no-repair(–HDR) control.

a. Transfer cell culture into sterile 15 mL tubes.
37. Wash cells with sterile water.

a. Centrifuge at 2500 g for 4 min and discard the supernatant.
b. Resuspend the pellet in 15 mL sterile water.
38. Resuspend in LiAc and aliquote cells per transformation.

a. Centrifuge at 2500 g for 4 min and discard the supernatant.
b. Resuspend the cell pellet in 0.1 M LiAc, using 1 mL per planned transformation (e.g., 3 mL total for 3 transformations).
c. For each transformation, transfer 1 mL of the LiAc cell suspension into a sterile 1.5 mL microcentrifuge tube. (e.g., 3 tubes for 3 transformations)
d. keep cell aliquots at room temperature while preparing transformation mixtures.
39. Prepare salmon sperm ssDNA.

a. Boil the salmon sperm ssDNA for 5 min.
b. Immediately cool the ssDNA on ice and keep it on ice until use.
40. Prepare the CRISPR transformation mixtures (350 µL total per transformation) in sterile 1.5 mL tubes.

a. Prepare one tube per transformation condition (e.g., for 3 transformations, prepare 3 separate tubes).
b. For the “CRISPR + HDR repair template” condition, add the following reagents in the indicated amounts:

| CRISPR + HDR repair template mixture |  |
| --- | --- |
| Reagents | Amount |
| Polyethylene Glycol (50% in H<sub>2</sub>O) | 240 μL |
| 1 M Lithium acetate | 36 μL |
| Salmon Sperm ssDNA | 25 μL |
| Plasmid contains Cas9 and sgRNA | 0.8-1 μg |
| HDR Repair oligo | 10 μg |
| Water | add to a final volume of 350 μL |

c. For the “CRISPR control (no repair template)” condition, add the following reagents:

| **CRISPR no-repair(–HDR) Control mixture** |  |
| --- | --- |
| Reagents | Amount |
| Polyethylene Glycol (50% in H<sub>2</sub>O) | 240 µL |
| 1 M Lithium acetate | 36 µL |
| Salmon Sperm ssDNA | 25 µL |
| Plasmid contains Cas9 and sgRNA | 0.8-1 µg |
| Water | add to a final volume of 350 µL |

41. Combine yeast cells with the transformation mixtures.

a. Centrifuge the 1 mL LiAc cell aliquots at 5000 g for 2 min.
b. Remove the supernatant completely.
c. Add 350 μL of the appropriate transformation mixture to each cell pellet.
d. Resuspend thoroughly by pipetting up and down until the pellet is fully dispersed.
42. Heat-shock the transformations.

a. Incubate the tubes at 42°C for 40 min in a water bath.
b. Centrifuge at 5000 g for 2 min and discard the transformation mixture completely.
43. Recover cells in YPD.

a. Add 1 mL YPD and resuspend the pellet.
b. Put caps on tubes and incubate at 30°C with shaking for 2-3 h (e.g., on a thermomixer).
44. Plate cells on selective medium.

a. Centrifuge at 5000 g for 2 min and discard the supernatant.
b. Add 150 μL sterile water and resuspend the pellet.
c. Spread the resuspended cells evenly onto the YPD + G418 (Geneticin) plates.
45. Incubate plates at 30°C until colonies appear (typically 2-3 days).
46. Assess transformation outcome. Expect many colonies on the “CRISPR + HDR repair template” plate and few colonies on the “no repair (–HDR) template” control plate (see Figure 9).

**Figure 9:**
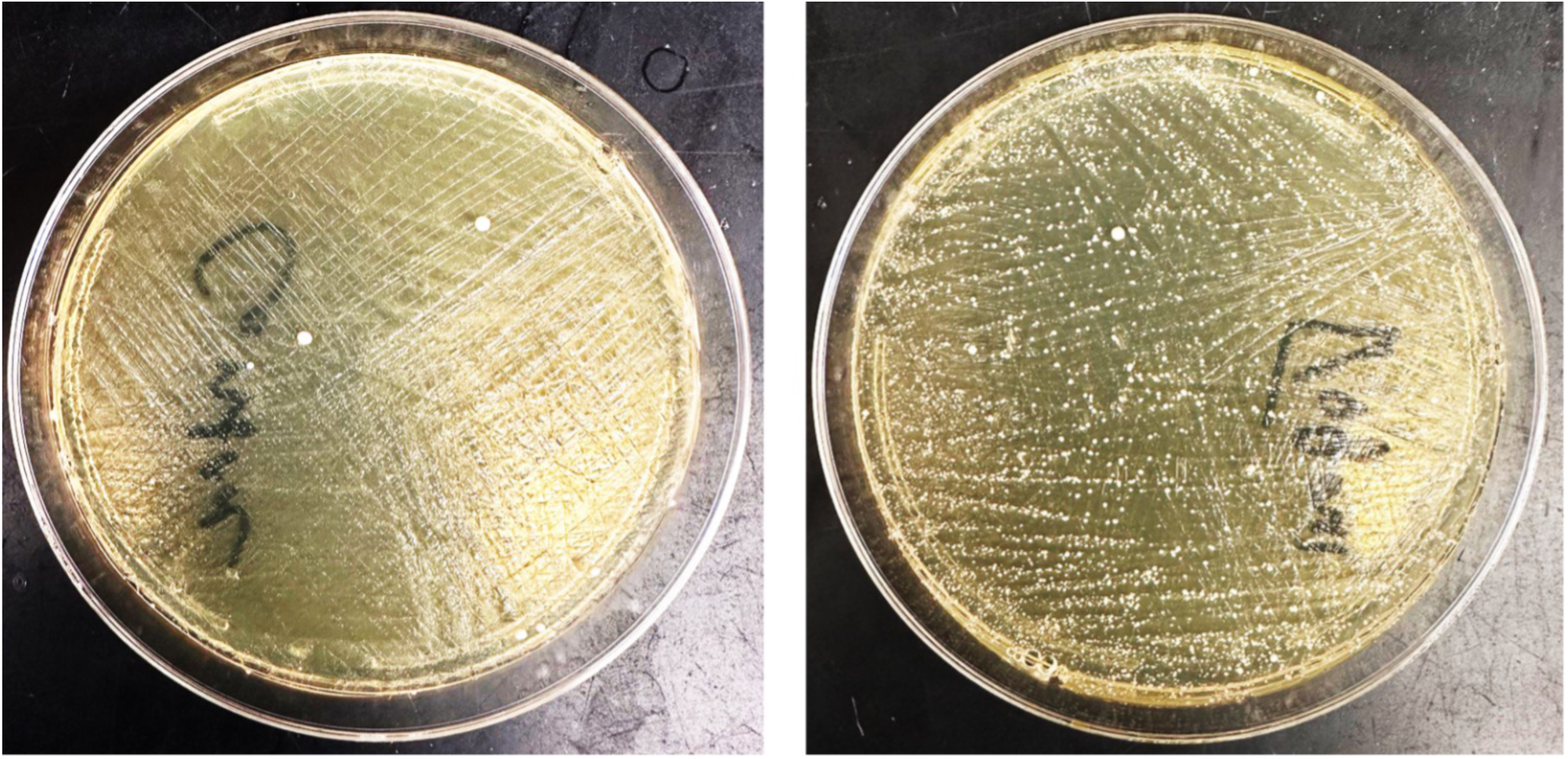
Yeast colonies on selective media plates following CRISPR-Cas9 transformation. (Right) Plate containing yeast cells transformed with the repair template, showing numerous colonies indicating successful homologous recombination and gene editing. (Left) Control plate with yeast cells transformed without the repair template, displaying significantly fewer colonies.

Note: A colony on the initial selection plate may contain a mixed population (edited and unedited cells). Restreaking yields single-cell–derived colonies.

47. Pick 8 colonies from the “CRISPR + HDR repair template”.
48. Streak each colony onto fresh YPD + G418 to isolate single colonies.

Note: To reduce the number of G418 plates used, divide each plate into 4-8 sectors and streak one colony per sector.

#### Verify genome editing

Timing: 2 days

This step identifies correctly edited yeast clones by PCR across the target locus and confirms the edit by Sanger sequencing.

CRITICAL: Use sterile tubes, tips, and solutions. Handle yeast colonies aseptically and work near a lit flame.

49. Pick single yeast colonies and prepare them for PCR.

a. Add 50 µL sterile water into 8 sterile PCR tubes
b. Pick 8 well-isolated colonies from the second YPD + G418 and resuspend each colony in 50 µL sterile water in PCR tubes.

CRITICAL: Store the PCR tubes containing the colony suspensions at 4°C until the sequencing results are available. After you confirm the correct edit, use the retained suspension to streak the corresponding clone onto plates for further culturing.

50. Set up a 50 μL colony PCR reaction for each clone using 5 μL of the cell suspension as the template.

a. Use primer pairs that flank the intended edit (i.e., primers anneal upstream of the left homology arm and downstream of the right homology arm).
b. Assemble the PCR mixture as follows:

| PCR reaction reagents |  |
| --- | --- |
| Reagents | Amount |
| 5× Reaction Buffer | 10 $\mu\text{L}$ |
| dNTP mix (2 mM each) | 1 $\mu\text{L}$ |
| Forward primer (10 $\mu\text{M}$) | 0.5 $\mu\text{L}$ |
| Reverse primer (10 $\mu\text{M}$) | 0.5 $\mu\text{L}$ |
| DNA template from yeast | 5 $\mu\text{L}$ |
| High-fidelity DNA polymerase | 0.5 $\mu\text{L}$ |
| Nuclease-free H<sub>2</sub>O | add to a final volume of 50 $\mu\text{L}$ |

51. Run the PCR program as follows

| PCR cycling conditions |  |  |  |
| --- | --- | --- | --- |
| Steps | Temperature | Time | cycle |
| Initial Denaturation | 95°C | 6 min<br>(for colony PCR) | 1 |
| Denaturation | 95°C | 30 sec | 34 |
| Annealing | 50-63°C (varied based on primers) | 30 sec |  |
| Extension | 72°C | Varied- 30 second per 1Kb size of the product |  |
| Final Extension | 72°C | 5 min | 1 |
| Hold | 4°C | Forever |  |

52. Analyze PCR products by agarose gel electrophoresis.

a. Confirm the expected amplicon size.
b. For deletions or insertions, confirm the expected size shift relative to an unedited control.

**Note:** If colony PCR yields multiple nonspecific bands (common for long amplicons), isolate genomic DNA and use 10 ng genomic DNA as the PCR template instead.

53. Submit PCR products for sequencing to confirm the edit at the nucleotide level.
54. After confirming correct edits, streak verified clones from the colony suspension (kept in water in PCR tube) onto YPD plates (non-selective) and grow without selection to promote loss of the CRISPR plasmid.

#### Expected outcomes

Following LiAc/PEG transformation and recovery, selection on YPD + G418 (Geneticin) enriches for yeast carrying the KanMX-marked CRISPR plasmid. After 2–3 days at 30°C, the “CRISPR + HDR repair template” condition typically yields many colonies, whereas the “no-repair (–HDR)” control yields few colonies (Figure 9), because Cas9 cutting is lethal unless repaired. This “many vs few” contrast is the initial readout that (i) transformation and selection worked and (ii) the chosen guide supports efficient cutting at the intended locus.

Downstream, the expected outcome from the workflow is a set of verified yeast clones carrying the desired edit. When screening 6–8 colonies by colony PCR using primers flanking the homology arms, correctly edited clones should produce an amplicon of the predicted size. Deletions and insertions should show a band shift relative to an unedited control, while point mutations may require confirmation by Sanger sequencing. For successful experiments, at least a subset of screened colonies is expected to be PCR-positive for the intended edit, and Sanger sequencing across the junction(s) should confirm the designed genotype, including any silent PAM-disrupting mutations included in the donor.

Finally, after outgrowth on non-selective medium (YPD), a fraction of edited clones should lose the KanMX-marked CRISPR plasmid; this can be evaluated by patching onto YPD and YPD+ G418 plate, where cured clones grow on YPD but fail to grow on G418.

#### Limitations

This protocol relies on SpCas9-induced double-strand breaks repaired by HDR, so success depends on guide cutting efficiency and HDR performance at the target locus. Editing can be less reliable for targets in repetitive or subtelomeric regions, for changes positioned farther from the cut site, or for edits that substantially reduce fitness (e.g., strong loss-of-function or essential-gene perturbations).

A second limitation is that the cloning workflow is tailored to a pML104-KanMX-sgRNAv2 backbone and In-Fusion Snap Assembly; using different plasmids or assembly methods may require re-optimizing primer overlaps and PCR conditions. Finally, accurate interpretation depends on effective G418 selection, which can vary by strain and can be confounded by pre-existing G418/KanMX resistance or weak antibiotic activity.

#### Troubleshooting

##### Problem1

No or weak full-length PCR band for the CRISPR plasmid (∼12 kb), or multiple bands or heavy smear in the whole-plasmid PCR (related to Steps 1-3). This can occur due to a suboptimal annealing temperature, poor primer design, overly high primer concentration, or poor template quality.

##### Potential solution

- Expand the annealing-temperature gradient and use the condition that yields the strongest single ∼12 kb band.
- Reduce primer concentration.
- Use fresh high-fidelity polymerase/buffer and a high-quality template plasmid.
- Try an alternative set of gRNA primers which includes sgRNA sequence.

##### Problem 2

Many *E. coli* colonies, but most plasmids are incorrect and/or contain carryover parental plasmid. This is commonly due to incomplete DpnI digestion of methylated parental plasmid or carryover of uncut template into the assembly/transformation (related to Steps 4-8 and 14-23).

##### Potential solution

- Confirm DpnI is active and extend digestion if needed; mix thoroughly before incubation.
- Purify the PCR product after DpnI digestion and avoid transferring any residual template stock.
- Sequence-verify the sgRNA region from multiple colonies before proceeding.

##### Problem 3

Few or no *E. coli* colonies after assembly/transformation (related to Steps 9-23). This can occur if the assembly reaction lacks correct overlaps, the competent cells are poor, or selection conditions are suboptimal.

##### Potential solution

- Recheck overlap design and DNA input amounts; use clean, quantified PCR product.
- Verify competent-cell performance with a control plasmid transformation.
- Confirm the plates/antibiotic match the plasmid selection used in your workflow.

##### Problem 4

Few or no colonies on both +HDR repair and –HDR repair plates (related to Steps 40-45). This can be due to low transformation efficiency. Alternatively, Cas9 cutting occurred, but HDR rescue failed, so even the +HDR condition cannot survive. Common causes include an insufficient or incorrect donor, a donor that does not disrupt the PAM or protospacer leading to re-cutting of the edited locus.

##### Potential solution

- Use log-phase cells (OD_600_ ∼0.6–1).
- Confirm all reagents is using in the yeast transformation (Polyethylene Glycol 50%, 1 M Lithium acetate and salmon sperm).
- Confirm 42°C heat shock for 40 min and 2–3 h recovery in YPD at 30°C before plating.
- If you suspect cutting without rescue confirm donor amount and integrity.
- Redesign the HDR donor so the repaired allele cannot be re-cut and make sure to mutate the PAM and/or introduce seed-region changes.

##### Problem 5

Many colonies on both +HDR repair and –HDR repair plates (related to Steps 40-45). Possible reasons include weak or absent cutting, so cells carrying the plasmid survive even without a donor (wrong guide sequence, poor guide activity, or poor expression). Alternatively, G418 selection may not be working, allowing plasmid-free cells to grow (wrong concentration, degraded antibiotic, or old/incorrect plates). Finally, the no-repair control may not have been truly –HDR (donor carried over or was added by mistake).

##### Potential solution

- Confirm the sgRNA plasmid is correct. Sequence the sgRNA-Cas9 plasmid to rule out primer mix-ups or incorrect guide insertion.
- Try a second sgRNA targeting the same locus which is often the fastest fix for no cutting.
- Verify G418 is selecting. Plate the same strain with no plasmid onto YPD + G418 in parallel. It should show no growth if selection is functioning. If plasmid-free cells grow well, remake fresh YPD + G418 and repeat the selection check.
- Confirm the –HDR control setup. Reconfirm donor DNA was truly omitted.

##### Problem 6

Colony numbers look right, but most colonies are wild type by colony PCR or sequencing (related to Steps 40-45). This suggests transformation and drug selection worked, but HDR or editing in survivors is inefficient. Possible causes include a suboptimal sgRNA, donor design that does not favor HDR at the intended junction, donor concentration or quality issues.

##### Potential solution

- Test a second sgRNA preferably closer to the edit site
- Ensure the HDR template blocks re-cutting (PAM/seed disruption), especially for point mutations and tags.
- Use an HDR template with longer homology arms.
- Increase screening depth: if the plate phenotype looks correct but the edit rate is modest, pick and test more colonies from the +HDR plate before concluding the design failed.

## Resource availability

### Lead contact

Further information and requests for resources and reagents should be directed to and will be fulfilled by the lead contact, **Brian D. Strahl**.

### Technical contact

Technical questions on execution of this protocol should be directed to and will be answered by the technical contact, **Hosein Rostamian**.

### Materials availability

The pML104-KanMX-sgRNAv2 plasmid generated in this study has been deposited to Addgene (plasmid #254712) and will be made available through Addgene upon publication.

### Data and code availability

This protocol did not generate or analyze any unique datasets or code. No custom code was used in this study.

## Acknowledgments

This work was supported by the National Institute for General Medical Sciences (NIGMS) under award R35GM126900 to B.D.S. and R35GM119518 to D. G. I.

## Author contributions

Hosein Rostamian developed and optimized the protocol, performed experiments, analyzed results, and wrote the original draft of the manuscript. Ethan W. Madden, Frank D. Kaplan, and Riley Kim contributed to protocol development, experimental validation/optimization, and editing of the manuscript. Daniel G. Isom contributed to conceptualization and plasmid generation. Brian D. Strahl supervised the study, contributed to conceptualization and interpretation, and edited the manuscript. All authors reviewed and approved the final version of the manuscript.

## Declaration of interests

B.D.S. is a co-founder of EpiCypher, Inc.

## Notes

### Competing Interest Statement

Brian Strahl is a co-founder of EpiCypher.

